# HICL table can manipulate all proteins in human complete proteome

**DOI:** 10.1101/093971

**Authors:** Zhenhua Xie

## Abstract

**Background:** The data of human complete proteome in the databases of Universal Protein Resource (UniProt) or National Center for Biotechnology Information(NCBI) were disorderly organized and hardly handled by an ordinary biologist.

**Results:** The HICL table enable an ordinary biologist efficiently to handle the human complete proteome with 67911 entries, to get an overview on the distribution of the physicochemical features of all proteins in the human complete proteome, to perceive the details of the distribution patterns of the physicochemical features in some protein family members and protein variants, to find some particular proteins.

Moreover, two discoveries were made via the HICL table: (1) The amino aicds(Asp,Glu) have symmetrical trend of the distributions versus pI, but the amino aicds(Arg, Lys) have local asymmetrical trend of the distributions versus pI in human complete proteome. (2) Protein sequence, besides amino acid properties, can in theory influence the modal distribution of protein isoelectric points.

**Conclusion:** I has created the HICL table as a robust tool for orderly managing 67911 proteins in human complete proteome by their physicochemical features, the names and sequences. Any proteins with the particular physicochemical features can be screened out from the human complete proteome via the HICL table. In addition, the unbalanced distribution of the amino aicds(Arg, Lys) in high pI proteins of human complete proteome and the effect of protein sequence on modal distribution of protein isoelectric points have been discovered through the HICL table.

## Background

A complete proteome is a group of proteins expressed by a <u>genome</u> completely sequenced. Sequences of the proteins in a complete proteome can be translated from all protein coding genes of a genome completely sequenced[1,2,3]. The availability of several thousand complete proteomes for the fully sequenced organisms has enabled us to decipher the evolutionary history of species through global comparative analyses[4,5]. However, it has been a challenging task for an ordinary biologist to handle a complete proteome with the massive amounts of entries. A coordinate system of a complete proteome should be developed for handling a complete proteome efficiently.

Mass spectrometry (MS)-based shotgun method is extremely powerful to analysize proteomes. Such a strategy relies heavily on the databases of complete proteomes[6,7]. In addition, 2D-PAGE approach has some its limitations: it can hardly separate very acidic, basic, small, large and hydrophobic proteins[8]. Therefore, organizing complete proteomes by physicochemical features of the protein sequences has become a strong need for the development of proteomics.

In order to interprete the biological functions of the many proteins in complete proteomes, sequence-sequence similarity or sequence-structure similarity play a critical role in predicting a possible function for a new sequence[9,10]. But these methods do not function properly when clear sequence or structural similarities do not exist as in case of far divergent evolution where sequence identities are below 25%[11]. Morover, not all homologous proteins have analogous functions. Some proteins have many shared domains, but they have different functions[12]. After all, because only ninety percent of proteins in the human complete proteome can matched at least one of 5494 manually curated Pfam-A families[13], the classification system based on sequence-sequence similarity of proteins is not a complete and user-friendly classification system. Therefore, the sequence-independent physicochemical features of proteins could be chosen as parameters to handle a complete proteome.

Proteins can be broken down into their constituent amino acids (AAs). Hydrophobicity, isoelectric point(pI), sequence length and molecular weight of a protein are independent of the sequence order information and only dependent of the numbers of amino acid composition (AAC) of the protein, so these physicochemical features have been designated as AAC-derived physicochemical features. The values of these features can be extracted simplely from a linear amino acid sequence. AAC and AAC-derived physicochemical features are powerful features that can predict protein-protein interactions, structural and functional classes of proteins and subcellular locations[14-17].

Excel is widely used by biologists for data manipulation [18]. In this study, like geographical coordinate system that uses degrees of latitude, longitude and altitude to illustrate a location on the earth’s surface, the values of AAC and AAC-derived physicochemical features had been chosen as multidimensional quantitative coordinates to locate all proteins in the human complete proteome. The values of intrinsic physicochemical features, ID numbers, names and Met-truncated sequences of all proteins in the human complete proteome had been organized as data matrix that was imported into Microsoft Excel(2007) to generate a excel table for manipulating of all protens in human complete proteome.

## Results and discussion

### The organization of the data in the HICL excel table

The HICL table was organized in 67912 rows and 28 columns and contains a header row and column for cell referencing. The numbers as showed in the header column represent the sequential numbers of the list of entries in the human complete proteome in FAST format, and the titles of columns in the header row use NO, Ala, Cys, Asp, Glu, Phe, Gly, His, Ile, Lys, Leu, Met, Asn, Pro, Gln, Arg, Ser, Thr, Val, Trp, Tyr, SL, MW, pI, HP, Annot1, Annot2, MTS to indicate the sequential number, the values of AAC, sequence length, molecular weight, pI, hydrophobicity, annotation and the corresponding Met-truncated sequence(MTS) of a protein. The all relative information of a protein was inserted in a corresponding row. The different parameters, annotations and MTSs of all proteins were respectively inserted in different corresponding columns for quickly manipulating the data of all protens in the human complete proteome. The annotation was devided to the protein name and ID number with the name abbreviation and -HUMAN for conveniently sorting all protens alphabetically according the names. All the names and ID numbers with the name abbreviations and -HUMAN were respectively inserted in the Annot1 and Annot2 columns.

Like geographical coordinate system that uses degrees of latitude, longitude and altitude to illustrate a location on the earth’s surface, using the value of AAC, sequence length, molecular weight, pI and hydrophobicity as numerical coordinates, a proteome coordinate system has been developed in the excel table to describe the location of every protein of the human complete proteome. Based on the tool of the Remove Duplicates section of the Data tab, 71 rows were detected as redundant rows in total 67911 rows according to the values of either AAC and sequence length or all physicochemical features in the data of the HICL table. Therefore, The values of AAC and sequence length can provide fundamental 21-dimentional coordinate system to locate all proteins. The values of molecular weight, pI and hydrophobicity derived originally from the values of AAC and sequence length can not provide additional information for locating the proteins, but as derived coordinates, they have crucial role for sorting, grouping and searching of all protens in the human complete proteome.

The data of all proteins in the human complete proteome have been organized as the data matrix in the HICL table, and the data matrix can be reorganized by the values of physicochemical features, names or the Met-truncated sequences.

### Sorting all proteins of the human complete proteome by physicochemical features

The sorted HICL table can illustrate both overview and detail of the distribution of AAC, sequence length, molecular weight, pI and hydrophobicity of all proteins. The all values of the every column in the HICL table were divided into five groups according to corresponding grouping criteria. Detail of the grouping criteria was illustrated in the table 1. The distributions of the all proteins in every numerical column of the HICL table were demonstrated in the table 2.

**Table1:** The values and ranges of the grouping criteria for every feature

| column | groupI | groupII | groupIII | groupIV | groupV |
| --- | --- | --- | --- | --- | --- |
| AAC | 0.0-<0.05 | 0.05-<0.1 | 0.1-<0.15 | 0.15%-<0.20 | ≥0.20 |
| SL | 0-<200 | 200-<500 | 500-<1000 | 1000-<2000 | ≥ 2000 |
| MW | 0-<23kd | 23kd -<57.5kd | 57.5kd -<115kd | 115kd -<230kd | ≥230kd |
| pI | 0-<4.0 | 4.0-<6.0 | 6.0-<8.0 | 8.0-<10.0 | ≥10.0 |
| HP | -<-1.0 | -1.0-< -0.5 | -0.5 -< 0.0 | 0.0-< 0.5 | ≥0.5 |

**The table 2:** The distributions of all proteins in the human complete proteome.

| feature | groupI | groupII | groupIII | groupIV | groupV |
| --- | --- | --- | --- | --- | --- |
| Ala | 15993 | 40795 | 9427 | 1344 | 352 |
| Cys | 61961 | 5218 | 543 | 74 | 115 |
| Asp | 40171 | 26228 | 1292 | 170 | 50 |
| Glu | 19625 | 38162 | 8435 | 1345 | 344 |
| Phe | 52084 | 15021 | 726 | 60 | 20 |
| Gly | 19763 | 39264 | 7437 | 1004 | 443 |
| His | 62991 | 4740 | 158 | 17 | 5 |
| Ile | 44232 | 22479 | 1115 | 72 | 13 |
| Lys | 30640 | 31115 | 5209 | 730 | 217 |
| Leu | 4359 | 29812 | 27788 | 5054 | 898 |
| Met | 66230 | 1601 | 69 | 9 | 2 |
| Asn | 55168 | 12312 | 385 | 39 | 7 |
| Pro | 27881 | 31382 | 6819 | 1364 | 465 |
| Gln | 41510 | 24404 | 1716 | 205 | 76 |
| Arg | 26816 | 35489 | 4681 | 722 | 203 |
| Ser | 9285 | 42439 | 13399 | 2172 | 616 |
| Thr | 32736 | 33031 | 1809 | 237 | 98 |
| Val | 22815 | 41142 | 3660 | 245 | 49 |
| Trp | 66602 | 1250 | 51 | 5 | 3 |
| Tyr | 62174 | 5513 | 175 | 28 | 21 |
| SL | 35926 | 19119 | 9045 | 3106 | 715 |
| MW | 36762 | 18872 | 8734 | 2887 | 656 |
| pI | 584 | 22680 | 15630 | 23892 | 5125 |
| HP | 4678 | 21003 | 31205 | 8388 | 2637 |

Lysine-rich proteins have nutritive and commercial value to establish transgenic lines of cereals with high lysine content of grains [20]. It was reported that down-regulation of cysteine-rich proteins and down-regulation of methionine-rich proteins can be respectively adopted by Escherichia coli and Synechocystis to sulfur deprivation[15]. Encoded by short open reading frames (sORF), small proteins take part in the developmental processes of plant and animal[21,22]. However, there has been no protocol to search any amino acid-rich proteins and small proteins in a complete proteome by now.

The HICL table integrates the the every protein name, ID number with the name abbreviation and -HUMAN and its MTS with its intrinsic values together, so it enables an ordinary biologist easely to make largescale analysis of the data, to perceive the details of the distribution patterns in the data, and to find all very acidic, basic, small, large and hydrophobic, highly cysteine-rich, highly aspartic acid-rich, highly glutamic acid-rich, highly lysine-rich, highly arginine-rich, highly serine-rich, highly threonine-rich, highly tryptophan-rich proteins and some other particular proteins in the human complete proteome. In addition, Ig heavy chain V-I region ND (Fragments)(|P01744|), Mucin-16(|Q8WXI7|), Mucin-19(|Q7Z5P9|) and Mucin-3A(|Q02505|) have the zero values of molecular weight and pI in the HICL table, because their sequences contain ambiguous amino acid character (X).

Any proteins with the values in the selected ranges of the physicochemical features can be screened out from the human complete proteome in the multi-sorted HICL table, for example, small-very acidic proteins, small-basic proteins, small-very acidic-hydrophobic proteins and small-basic-hydrophobic proteins. All these specific proteins will give us an explicit information to design a better preparation and separation protocol in human proteomics research. In adition, all Met-truncated sequences of the proteins with the values in the selected ranges of the physicochemical features can be copy from the multi-sorted HICL table for further analysis.

### Sorting all proteins of the human complete proteome by the names

The rearrangement of the data in the HICL table can be accomplished by alpha-betically sorting of the Annot1 column. Some proteins which names are beginning with the same letter are grouped together; and within that grouping all proteins which names are beginning with the same two-letter sequence are grouped together; and so on. The rearranged data can show all proteins of the human complete proteome in alphabetical order based on their names. Some protein family members or protein variants can be generally grouped together in clusters, because the initial alphabets of their names are identical. This sorted table enable an ordinary biologist quickly to visualize and find the details of the distribution of physicochemical features in some protein family members and protein variants. For example, the majority of total 419 members of olfactory receptor family possess the high hydrophobicity values ranged from 0.5085 to 1.089, except for Olfactory receptor 4N2 (Fragment) (-0.2999), Olfactory receptor 2T12 (0.4367), Olfactory receptor 1L1(0.4496), Olfactory receptor 8S1(0.462), Olfactory receptor 2T33(0.4683).

There are the groups of the proteins which names are beginning with ankyrin, cyclin, cysteine-rich, DDB1, DNA, F-box, histone, nucleolar, olfactory, PDZ, transmembrane, zinc finger, etc. For example, figure1 shows the cluster of protein family members and protein variants of alpha-2,8-sialyltransferase 8 in the HICL table sorted by the names.

**Figure1:**
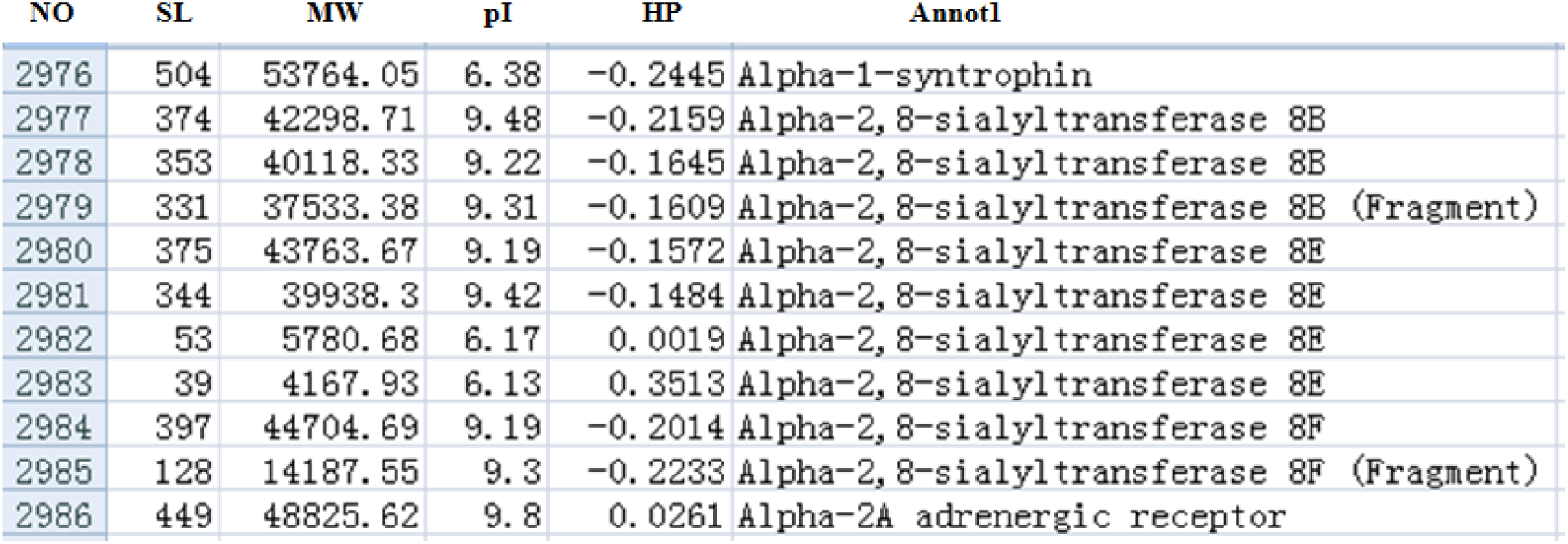
The cluster of alpha-2,8-sialyltransferase 8 family members and variants in the HICL table sorted by the names

The total number of the 20,687 protein-coding genes were predicted from the human genome[23], therefore, the set of 67911 entries in complete human proteome must contain many protein variants. So far, there has been no protocol to cluster all protein variants together in a complete proteome. By means of sorting all proteins of the human complete proteome by the names, all annotated protein variants can be comprehensively organized in clusters. so it enables an ordinary biologist easely to observe and perceive the all clusters of the protein variants in the sorted HICL table.

### Sorting all proteins of the human complete proteome by the Met-truncated sequences

The HICL table can be converted by alphabetically sorting of the MTS column. Some Met-truncated sequences are beginning with the same letter are grouped together; and within that grouping all Met-truncated sequences are beginning with the same two-letter sequence are grouped together; and so on. The rearranged data can show all proteins of the human complete proteome in alphabetical order based on their Met-truncated sequences and enable an ordinary biologist to make largescale analysis of the N-terminal amino acid sequences in the human complete proteome. Interestingly, some protein family members or protein variants can be usually grouped together in clusters, because their N-terminal amino acid sequences are identical. Some protein family members and protein variants have different N-terminal amino acid sequences, so they may take mixed pattern of dispersed and aggregated distributions in the HICL table sorted by the Met-truncated sequences.

Most neighbouring different proteins have the same two or three amino acid residues in their N-terminal amino acid sequences. Thus, like in some prokaryote proteome projects[24,25], N-terminal amino acid sequences of the human complete proteome have sufficient specificity for protein identification in human proteome projects.

### Searching of all protens in the human complete proteome by query sequences or the names

The data of the HICL table can be quickly searched by the text of query sequences, the names or part of name. The peptides from Mass Spectrometry (MS)-based peptide sequencing can be as query sequences to search the precursor sequences in the HICL table. Using the words of ankyrin, cyclin, cysteine-rich, DDB1, DNA, F-box, histone, nucleolar, olfactory, PDZ, receptor, transmembrane, zinc finger, etc, some protein groups can be quickly identified and located by searching the alphabetically sorted data in the HICL table.

In comparison with the HICL table sorted by protein names, the HICL table sorted by the Met-truncated sequences has no alphabetical order in protein names. The difference of N-terminal amino acid sequences of protein family members and protein variants can be estimated by their distribution in the HICL table sorted by the Met-truncated sequences. Therefore, the function of searching is critical to reveal the distribution of protein family members and protein variants in the HICL table sorted by the Met-truncated sequences. For example, figure2 shows the result of the searching by the text of alpha-2,8-sialyltransferase 8 and demonstrates the distribution pattern of protein family members and protein variants of alpha-2,8-sialyltransferase 8 in the HICL table sorted by the Met-truncated sequences.

**Figure2:**
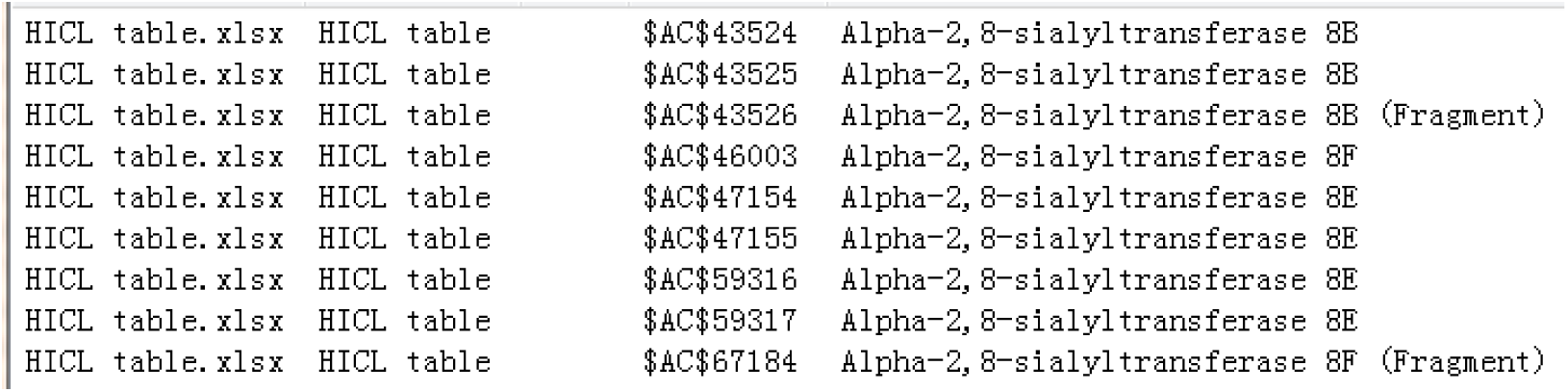
The distribution of alpha-2,8-sialyltransferase 8 family members and variants in the HICL table sorted by the M-truncated sequences

### Illustrating the physicochemical maps of the human complete proteome

The data matrix in the HICL table contains multidimensional quantitative coordinates to locate all proteins in the human complete proteome. The all proteins of the human complete proteome can create a map in the multidimentional space. This map can be described as the physichemical structure of the human complete proteome, but can not be directly demonstrated for us. This map can be projected into any coordinate or two-dimensional space to generate an image to visualize the distribution patterns of one or two physicochemical features in the human complete proteome. The pI distribution in human complete proteome show a bimodal distribution(figure3). The distributions of individual amino acid versus sequence length were demonstrated in the additional file named as “distributions versus sequence length’’. The maps in Figure4 were selected from the additional files to show the distributions of the amino aicds (Cys, Leu, Lys,Pro) versus sequence length. The distributions of the amino aicds (Asp,Glu,Arg, Lys,His, Cys,Tyr) with an ionizable side-chain versus pI were all demonstrated in figure5.

**Figure3:**
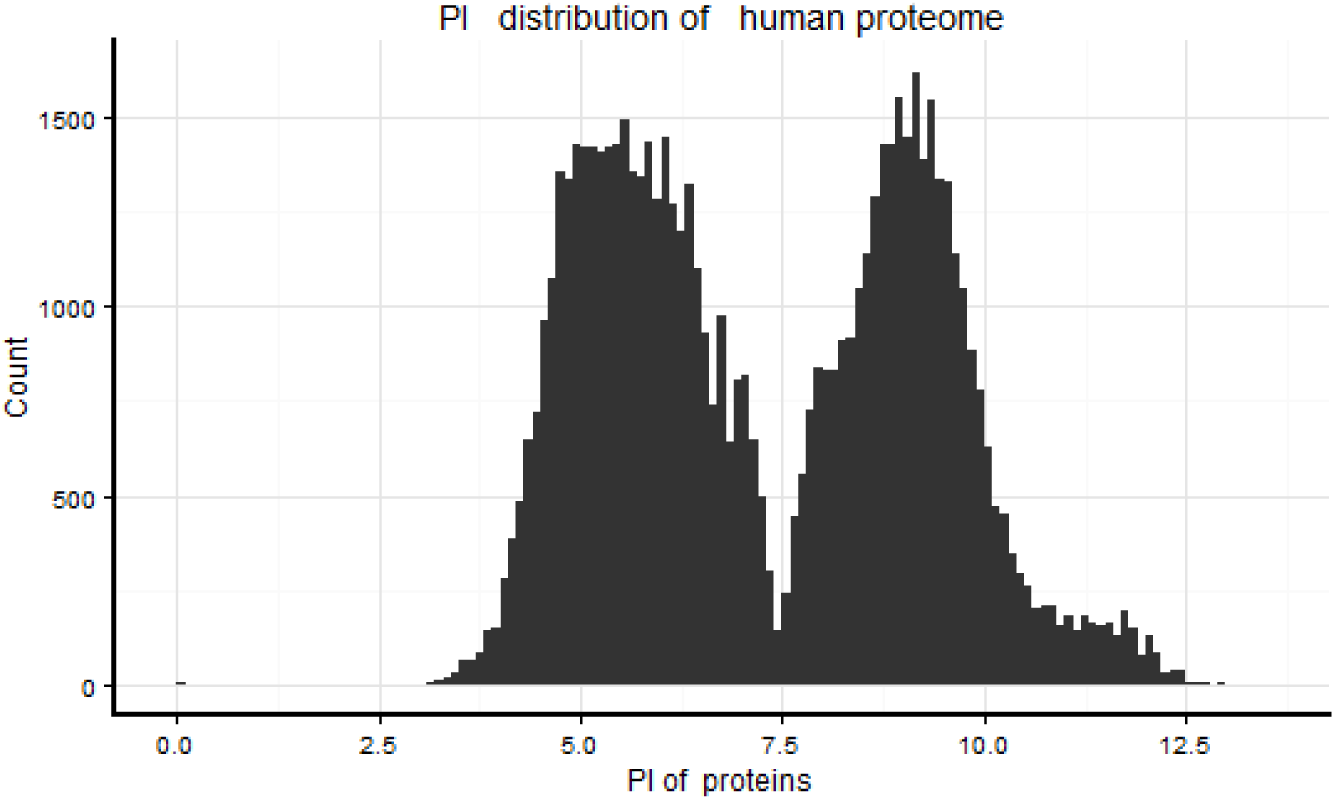
The pI distribution in human complete proteome

**Figure4:**
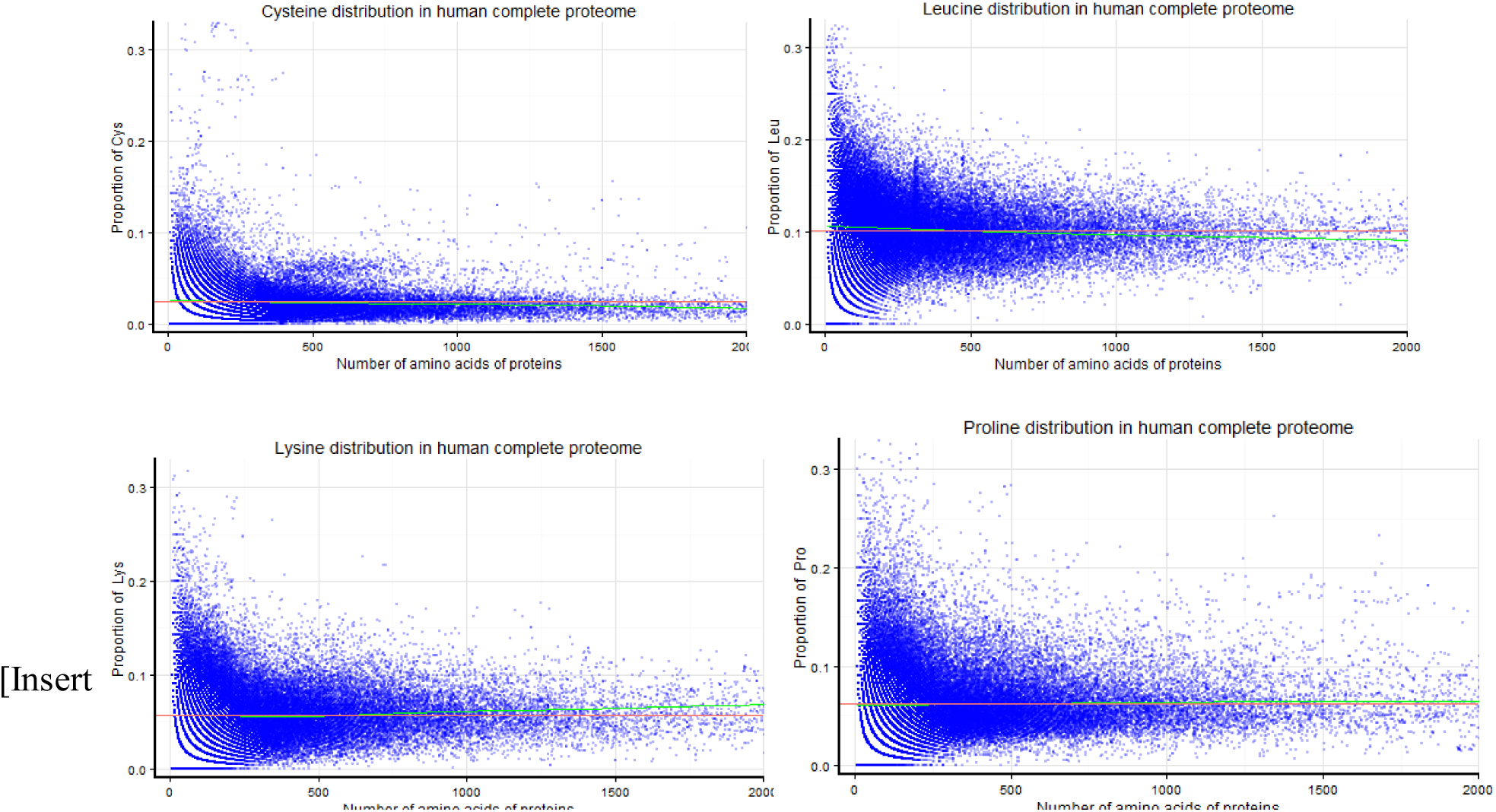
The distributions of the amino aicds (Cys, Leu, Lys,Pro) versus sequence length

**Figure5:**
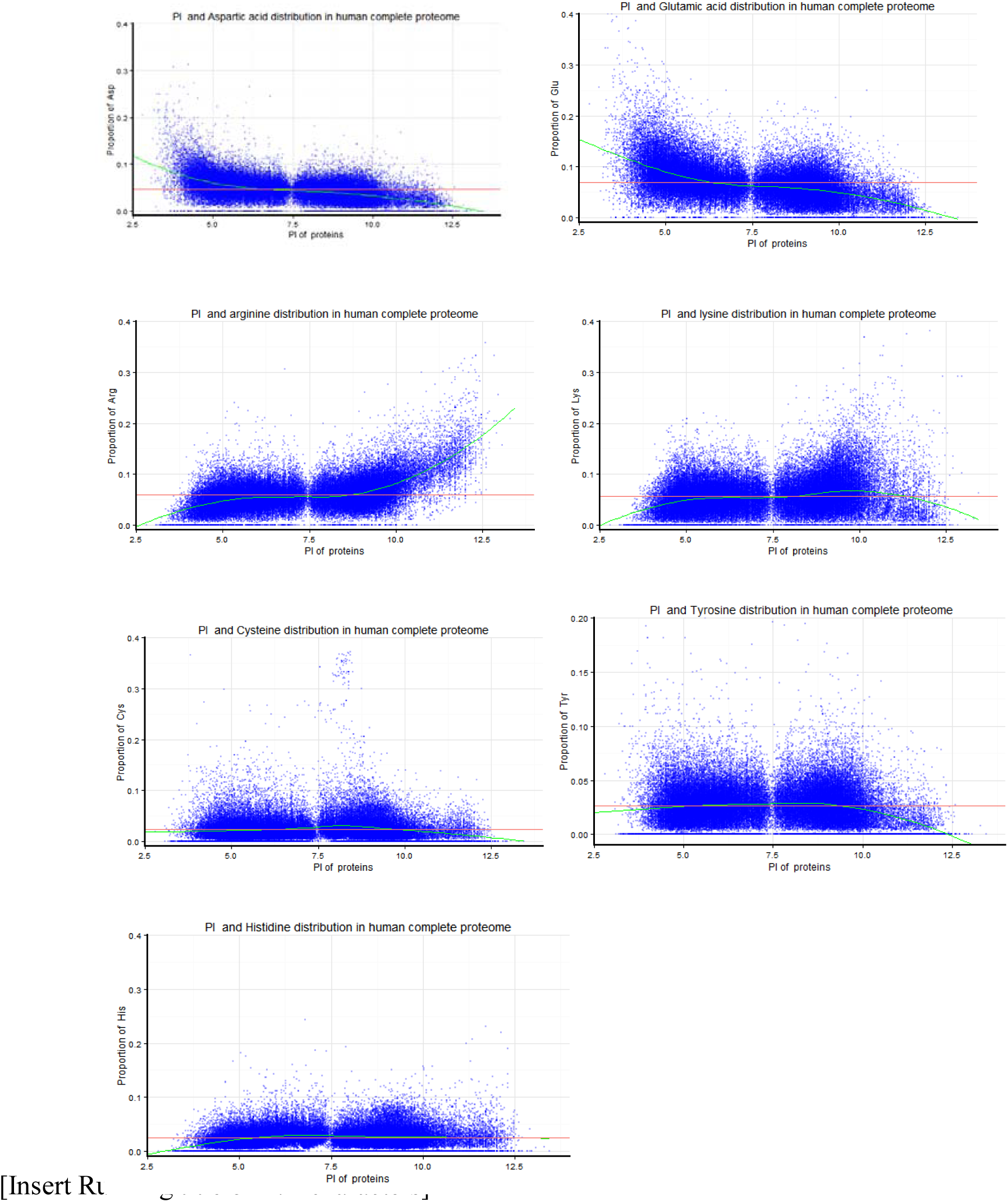
The distributions of the amino aicds (Asp,Glu,Arg, Lys,His, Cys,Tyr) versus pI

According to this bimodal distribution(figure3), normal acid-base state from 7.35 to 7.45 of pH values in human blood is beneficial for the stability of all human proteins to avoid the aggregation of protein at pI as best as possible. In the maps of the distributions of individual amino acid versus sequence length, the green loess curves extends tightly around the corresponding red average line in the range of about sequence length≤ 1000 and then gradually deviates from the corresponding red average line in the range of about sequence length > 1000. It means that any subgroup proteins in the range of about sequence length≤ 1000, for example, small proteins, has almost as same as the average values of amino acid composition in human complete proteome. With weak acid-base groups and low-abundance in the human complete proteome, the amino aicds(His, Cys, Tyr) have L-shaped trends of distributions versus pI, but the trends of the amino aicds(Cys,Tyr) with weak acid groups decline in the range of high pI and the trend of the amino aicd(His) with weak base group declines in the range of low pI. With strong acid-base groups and high-abundance in the human complete proteome, the amino aicds(Asp,Glu,Arg) have Sshaped trends of distributions versus pI, but the trend of the amino aicd(Lys) declines in the range of low pI and high pI and have C-shaped trends of distributions versus pI. In a word, the amino aicds(Asp,Glu) have symmetrical trend of the distributions versus pI, but the amino aicds(Arg, Lys) have local asymmetrical trend of the distributions versus pI in human complete proteome.

### Creating a particular fusion proteome via the HICL table

By numerically sorting the pI column in either ascending or descending order,all Met-truncated sequences in MTS column were respectively copied and pasted into A and B columns in sheet1, and then sheet1 was saved as a txt document named as fusion-proteome1. The values of isoelectric point(pI) of all fusion proteins in fusion-proteome1 had been computed using Compute pI/Mw tool (http://web.expasy.org/compute_pi/). The pI distribution in fusion-proteome1 has been illustrated in figure6.

By comparison between figure3 and figure6, it could be asserted that protein sequence, besides amino acid properties, can influence the modal distribution of protein isoelectric points and result in the maldistribution of protein isoelectric points in particular fusion proteome. This result supplements Georg F. Weiller’s idea: “The modal distribution of protein isoelectric points reflects amino acid properties rather than sequence evolution”[26].

**Figure6:**
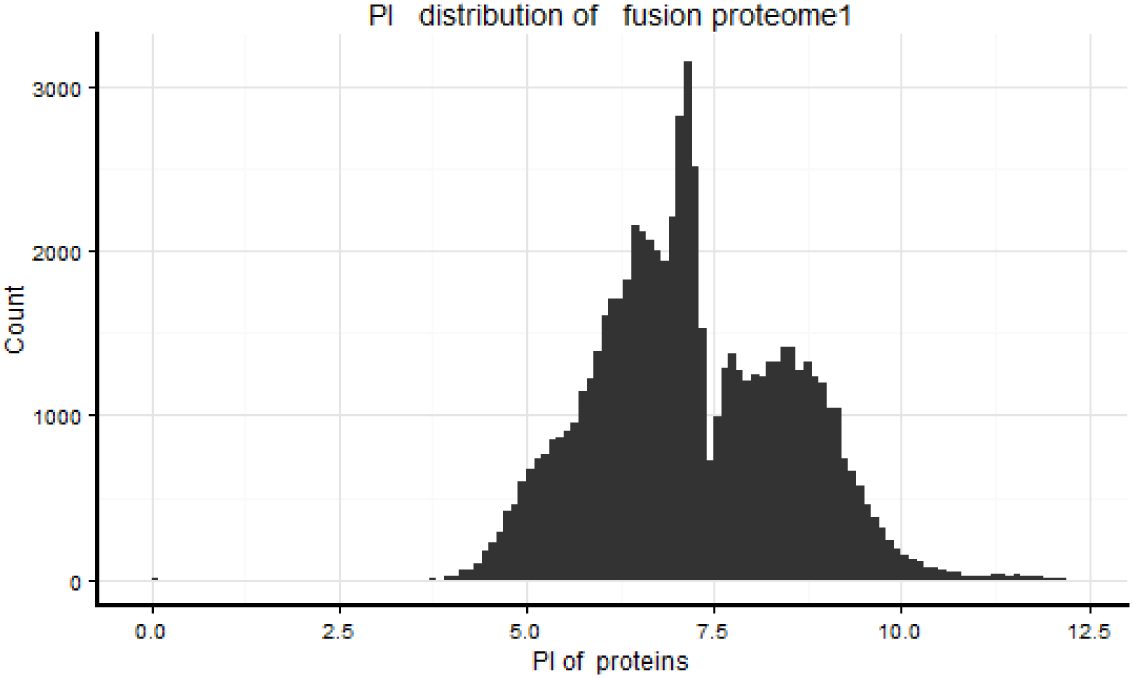
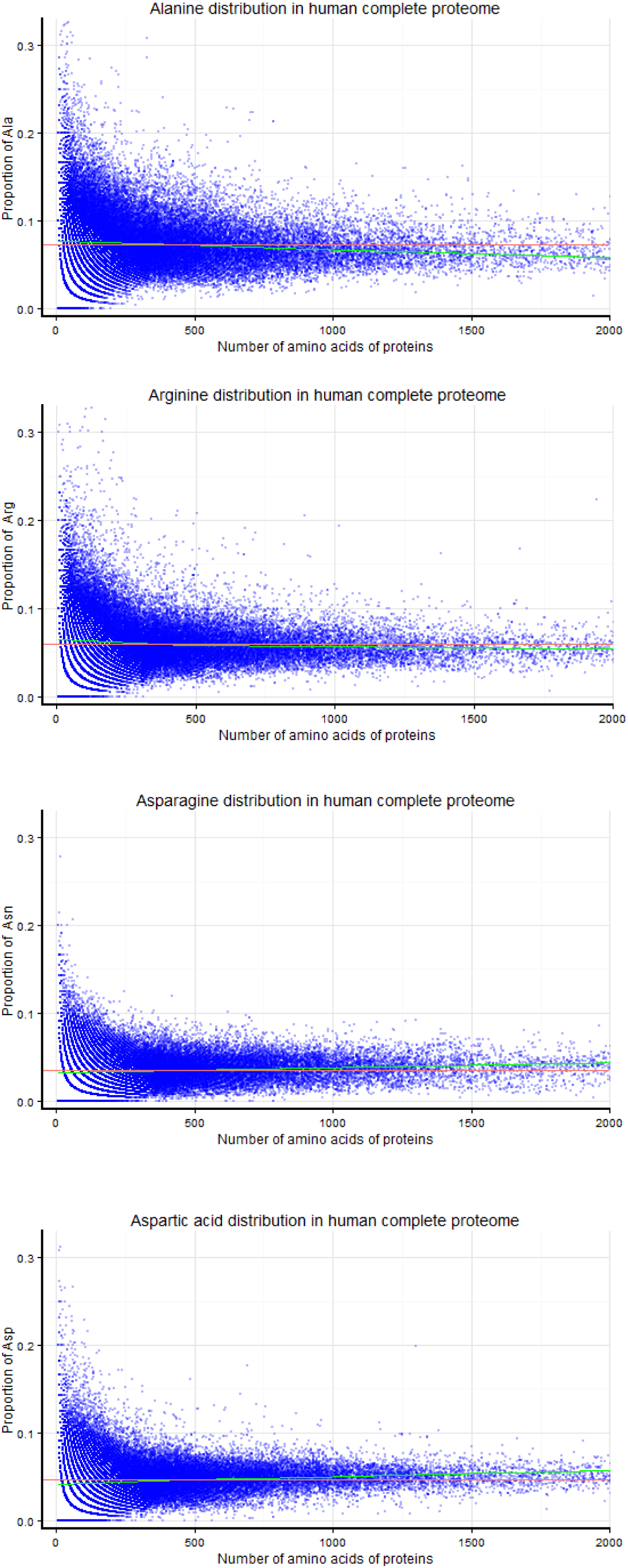
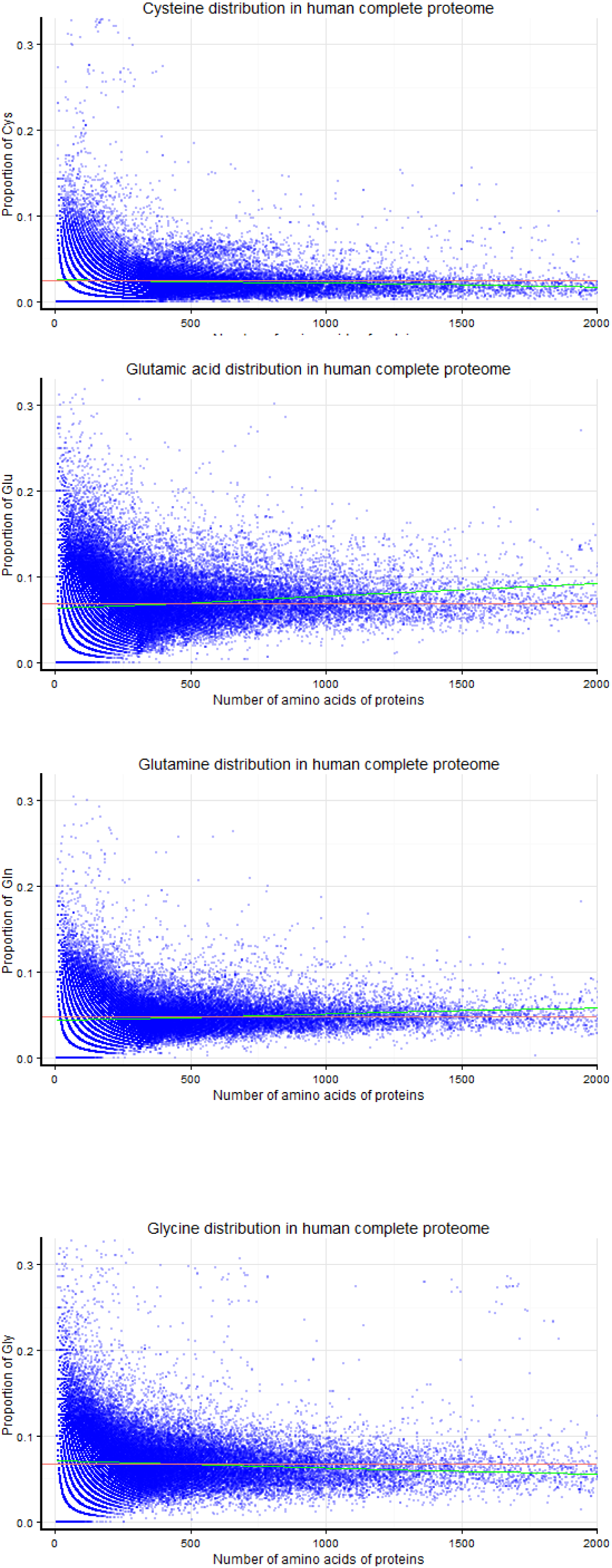
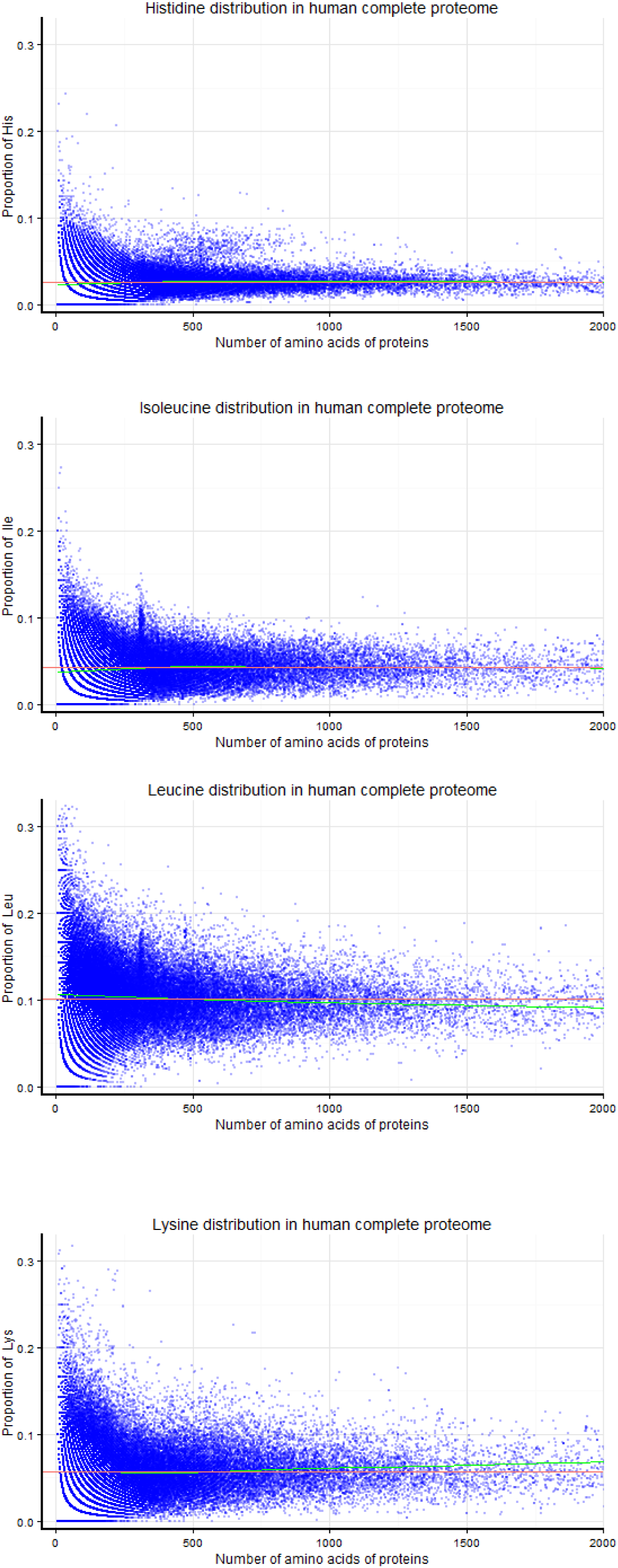
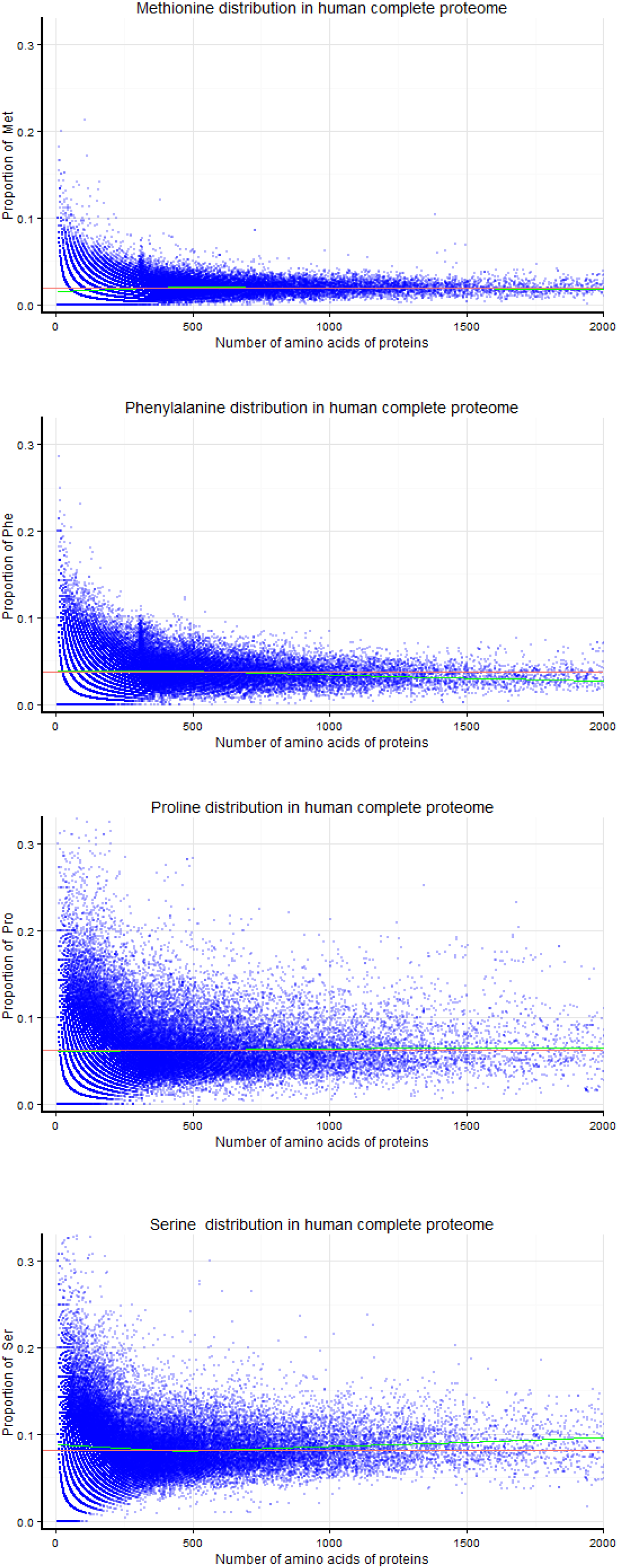
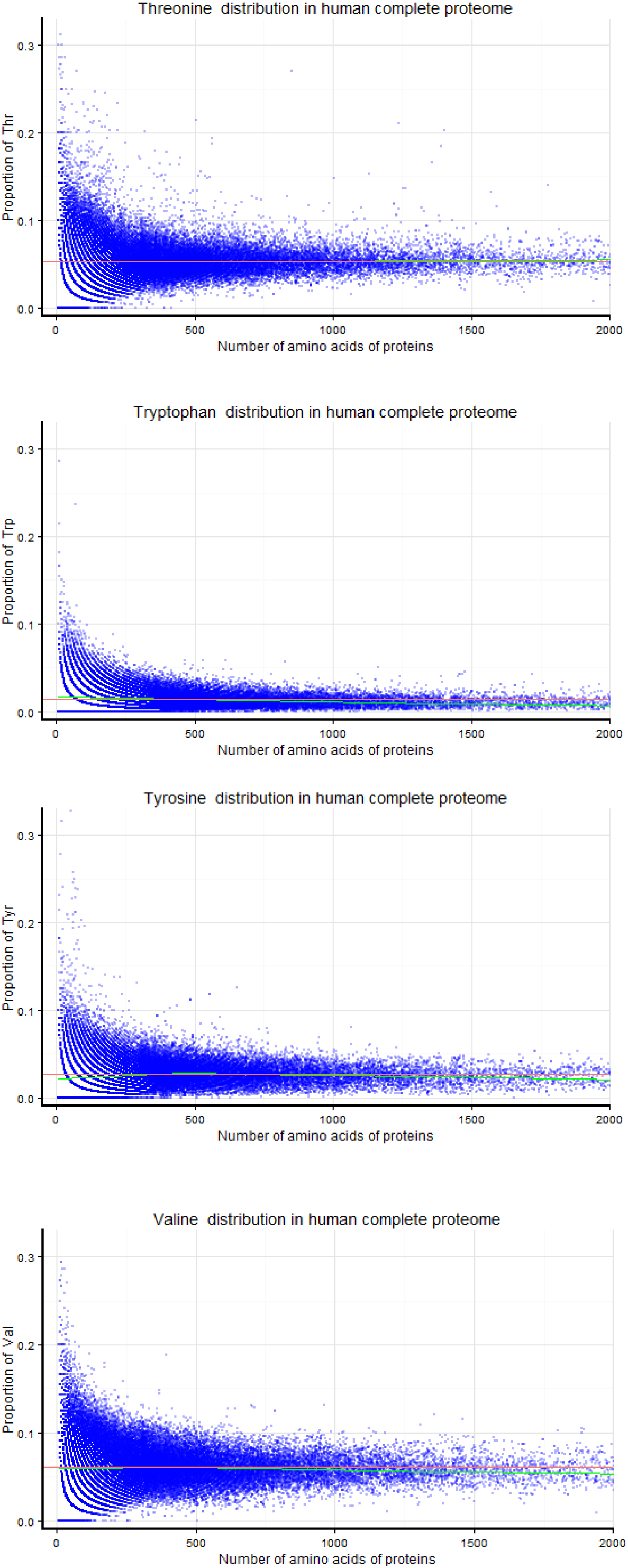
The pI distribution in fusion-proteome1

## Conclusions

This HICL table can be orderly reorganized by numerically or alphabetically sorting any colume. Any proteins with the values in the selected ranges of the physicochemical features can be screened out from the human complete proteome in the multi-sorted HICL table, and all very acidic, basic, small, large, hydrophobic, highly aspartic acid-rich, highly glutamic acid-rich, highly lysine-rich, highly arginine-rich and some other particular proteins can be easily found in the human complete proteome. Some protein family members or protein variants can be generally grouped together in clusters, because the initial alphabets of their names or their N-terminal amino acid sequences are identical. The data of the HICL table can be quickly searched by the text of query sequence, the name or part of name, so any protein family can be quickly identified and located by searching the alphabetically sorted data in the HICL table. Based on the data in the HICL table, the distribution patterns of any one or two physicochemical features in the human complete proteome could be visualized. A particular fusion proteome can be created via the HICL table for theoretical research of some feature in proteome.

The HICL table contains almost complete information of the set of human complete proteome entries downloaded from the Universal Protein Resource (UniProt) and the values of AAC, AAC-derived features of all proteins. Based on the integration of the data matrix and the functions of sorting and searching, an overview on the distribution of the physicochemical features of all proteins in the human complete proteome and the details of the distribution patterns of the physicochemical features in some protein family members and protein variants can be quickly illustrated in the HICL table. Therefore, the HICL table can be a robust tool for exploiting the human complete proteome and for exploring the feature of proteome. This method can be applied to the complete proteomes of other species.

Based on the data in the HICL table, the unbalanced distribution of the amino aicds(Arg, Lys) in high pI proteins of human complete proteome and the maldistribution of protein isoelectric points in particular fusion proteome have been discovered.

## Materials and Methods

### The establishment of HICL excel table

The set of human complete proteome entries in FASTA format had been downloaded from the Universal Protein Resource (UniProt) [1] and demonstrated in the additional file named as ‘‘human complete proteome’’. The original sequences of proteins in FASTA format had been transformed into the amino acid sequences of the proteins in plain-text format that were then converted into the Met-truncated sequences(MTSs) by eliminating the initial methionine. The abundances of amino acids in the MTSs can be calculated as the values of AAC. The MTSs and annotations of the human complete proteomes had been extracted from the set of human complete proteome entries by the R statistical programming language. The values of AAC, sequence length, molecular weight, isoelectric point(pI) and hydrophobicity of all MTSs in human complete proteome had been computed using the R statistical programming language, ProPAS software[19] and Compute pI/Mw tool (http://web.expasy.org/compute_pi/). The all values of the results with corresponding MTSs and annotations were imported into Microsoft Excel(2007) to generate a excel table for further operating analysis. This excel table has been designated as HICL.

### The manipulation of the HICL excel table

In Excel, the tool on the Remove Duplicates section of the Data tab can provide the function for removing all duplicate rows from the range of data and leaving only the first instance of each row. Excel displays a message confirming how many duplicates were removed and the number of unique values remaining, so the number of redundant rows of numeric values of physicochemical features in the data of the HICL table can be determined.

Using the tool on the Sort & Filter section of the Data tab, the data of a excel table can be reorganized by numerically or alphabetically sorting a specific column in either ascending or descending order (from the smallest to largest numeric value or from largest to the smallest numeric value) and ( A to Z or Z to A). The data of the HICL table can be quickly reorganized by respectively sorting any column.

The HICL table has its multidimensional quantitative coordinates, so the data in the HICL table can be grouped according to not only a coordinate but also their coordinates. After sorting by first physicochemical feature, the rows in which the values of first physicochemical feature are below or/and above criteria can be deleted. The deleted data of the HICL table continues to be sorted by second physicochemical feature, New reorganized data can show the details of the second physicochemical feature distribution of the proteins with the values in the selected range of first physicochemical feature. To continues these steps with other physicochemical feature, and so on, any proteins with the values in the selected ranges of the physicochemical features can be screened out from the human complete proteome.

Using the tool on the Find & Select section of the Home ribbon, the data of the HICL table can be searched by opening the Find and Replace dialog, pasting or typing the text of query sequence, the protein name or part of name that you are searching for in the "Find" box, and then Clicking "Find All" to generate a list of all rows that contain that text.

### The illustration of the distribution of physicochemical features in the human complete proteome

Based on the data in the HICL table, the map of the distribution of physicochemical features in human complete proteome had been illustrated by the R statistical programming language. In the maps g, the green loess curve and the red average line can clearly show the trend of an individual amino acid distribution.

## Ethics

Ethical approval was not applicable for this type of study.

## Consent to publish

Any consent to publish was not applicable for this study.

## Authors’ contributions

Zhenhua Xie did all works in this study and wrote the manuscript.

## Acknowledgements

The author would like to acknowledge the supports from Shenzhen Bureau of Science,Technology and Information (Grant No. JCYJ20140417115840267 and JCYJ20150518162154828).

## Additional files

Supplementary HICL excel table and the additional files ‘‘distributions versus sequence length’’and ‘‘human complete proteome’’ can be found online at‑‑‑‑‑‑‑

## Abbreviations

2D-PAGE: two-dimensional polyacrylamide gel electrophoresis
AAC: Amino acid composition
AAs: amino acids
Ala: Alanine
Annot1: Annotation1
Annot2: Annotation2
Arg: Arginine
Asp: Aspartic acid
Asn: Asparagine
Cys: Cysteine
DDB1: damage-specific DNA binding protein1
DNA: deoxyribonucleic acid
F-box: a protein structural motif of about 50 amino acids that mediates protein–protein interactions
Gln: Glutamine
Glu: Glutamic acid
Gly: Glycine
His: Histidine
HP: Hydrophobicity
ID: identification
Ile: Isoleucine
Leu: Leucine
Lys: Lysine
Met: Methionine
MS: Mass spectrometry
MTS: Met-truncated sequence(derived from full protein sequence by eliminating the initial methionine)
MW: Molecular weight
NCBI: National Center for Biotechnology Information
NO: Number
PDZ: a common structural domain of 80-90 amino-acids found in the signaling proteins of bacteria, yeast, plants, viruses[1] and animals
Phe: Phenylalanine
pI: Isoelectric point
Pfam: a large collection of protein families, each represented by multiple sequence alignments and hidden Markov models (HMMs)
Pro: Proline
Ser: Serine
SL: Sequence length
sORF: short open reading frames
Thr: Threonine
Trp: Tryptophan
Tyr: Tyrosine
UniProt: Universal Protein Resource
Val: Valine

